# Selection vs. integration task demands shape the similarity of information neural coding

**DOI:** 10.64898/2026.05.26.727806

**Authors:** Blanca Aguado-López, Ana F. Palenciano, María Ruz

**Author notes:** **Corresponding author:** Blanca Aguado-López.

## Abstract

Attention is a function that enables selection and integration of multiple sources of information. However, how these demands influence neural coding of information is not well understood. In this study we used EEG to examine how the selection vs. integration of stimuli shapes the content and geometry reflected on neural patterns, during both preparation and target processing. Participants performed a size judgement task in a cue-target paradigm that, depending on the block, required judging either the size of a selected item and ignoring the additional stimulus or integrating both to respond. Decoding analyses showed that under selection demands, categorical templates of the cued stimulus were active during preparation and target coding, contrasting with integration, where patterns linked to the cued category could be observed only during preparation. Notably, RSA suggested specific exemplar encoding during its processing, which was sustained also across the post-stimulus window during selection, yet was not observable under integration contexts. Our results also suggest that attentional demands shape the similarity between stimulus categories by increasing the distance between selected stimuli and distractors or by reducing it between to-be-integrated information. Overall, this study uncovers the dynamics of stimulus encoding under selection and integration demands, offering crucial advances to understand how top-down processes shape information representation in the human brain.

## 1. Introduction

Selective attention is a complex cognitive function that enables the *selection* of information by filtering unwanted data and prioritizing stimuli that are relevant to us (Driver, 2001). Additionally, attention facilitates the *integration* of information by focusing and combining multiple sources (Treisman & Gelade, 1980). While a substantial body of research has investigated selective attention of single stimuli (Aguado-López et al., 2024; Desimone & Duncan, 1995; Kaiser et al., 2016; Peñalver et al., 2023; Stokes et al., 2009) studies examining attention to multiple items remain scarce (Haynes et al., 2005; Kim et al., 2017; McMains & Somers, 2004; Morawetz et al., 2007; Treisman & Gelade, 1980). To advance this knowledge, the current study examined how selecting and integrating information about objects differentially impacts categorical and stimulus-specific neural coding.

One of the most influential accounts of selective attention is the biased competition model (Desimone & Duncan, 1995). This framework emphasizes the limited capacity of neural processing and the role of competition among sensory inputs for neural representation. Such competition is modulated by bottom-up and top-down factors. While bottom-up flow may favor salient stimuli, top-down mechanisms recruit prefrontal neurons to generate internal templates that bias competition toward goal-relevant information. Complementing this view, integration involves combining information across different sources and levels of representation. Traditionally, the Feature Integration Theory (Treisman & Gelade, 1980) proposes that attention is required to bind different features into a coherent representation of an individual object. However, attention may also be required when information from multiple distinct objects must be jointly considered to guide a behavioral decision (Duncan, 1984), a topic that has been much less explored.

A variety of experimental paradigms have been used to investigate these processes. Studies of selective attention commonly employ cues that direct attention toward specific stimuli or features while requiring participants to ignore others (Aguado-López et al., 2024; Desimone & Duncan, 1995; Peñalver et al., 2023; Stokes et al., 2009). A common finding is that interference arises when targets and distractors demand opposing responses. By comparison, studies on integration require either binding features from the same object for its location and identification as proposed by the Feature Integration Theory (Seymour et al., 2010; Treisman & Gelade, 1980) or attending to multiple stimuli or spatial locations. For example, Duncan (1984) showed that participants could make two judgments simultaneously when they concerned the same object, but performance was impaired when the judgments concerned different objects, highlighting the role of object-based attention. More recent studies have examined situations in which multiple spatially distinct stimuli are simultaneously task-relevant and their information must be jointly considered (Fagioli & Macaluso, 2016; Haynes et al., 2005; Kim et al., 2017; McMains & Somers, 2004; Morawetz et al., 2007). When stimuli are located across space, it is found that spatial attention may be divided across different spatial regions (McMains & Somers, 2004; Morawetz et al., 2007), although performance is worse when more than one object needs to be attended (Fagioli & Macaluso, 2016; Morawetz et al., 2007). Nonetheless, there is scarce literature comparing selection and integration demands when post-perceptual information of objects needs to be considered to generate a response.

At the neural level, selected target stimuli are processed with greater precision as reflected in enhanced neural responses in parietal, frontal and posterior cortices (Desimone & Duncan, 1995). Complementing these findings, oscillatory studies show that target selection is accompanied by increased parietal and midfrontal theta power (Haciahmet et al., 2021), highlighting the role of low-frequency dynamics in top-down attention. Building on this, analytical techniques such as multivariate pattern analysis (MVPA) have revealed that brain activity patterns encode attended stimuli or features, showing more robust representations of stimulus-specific characteristics (Kaiser et al., 2016; Reddy et al., 2009; Sheldon et al., 2021). These results have been interpreted as evidence for attentional templates: task-relevant representations that are maintained or activated during preparation and that specify information needed to guide subsequent stimulus processing (Desimone & Duncan, 1995). Supporting this view, it has been shown that such templates can be preactivated before stimulus onset, including shape (Stokes et al., 2009), category (Aguado-López et al., 2024; Esterman & Yantis, 2010; González-García et al., 2017; Peelen & Kastner, 2011; Peñalver et al., 2023) and spatial templates (González-García et al., 2016; Rajan et al., 2021). Altogether, these findings indicate that selective attention optimizes processing by enhancing and preconfiguring neural codes for relevant stimuli. Relatedly, attentional integration of multiple targets is found to increase activity in prefrontal and parietal cortices (Fagioli & Macaluso, 2016; Kim et al., 2017) and to enhance functional connectivity among visual areas (Haynes et al., 2005). When targets appear at different spatial locations simultaneously, visual cortex responses are selectively amplified for each attended position (McMains & Somers, 2004; Morawetz et al., 2007), suggesting that attentional resources can be split generating multiple spotlights. However, since most research has addressed either selection or integration demands independently, it is still unknown how these two requirements differ in preparatory attentional templates and evolve during actual stimulus perception.

The present study addresses this gap by examining whether and how the requirement to process one versus multiple task-relevant stimuli shapes the neural representations involved in their processing over time. To explore this, we collected electroencephalography (EEG) using a cue–target paradigm designed to keep the stimulus conditions comparable across tasks, while manipulating whether participants had to base their response on information from a single stimulus (selection condition) or from two stimuli (integration condition). In light of previous findings (Aguado-López et al., 2024; Esterman & Yantis, 2010; González-García et al., 2017; Peñalver et al., 2023), an intriguing hypothesis is that the coding geometry is tailored by attentional demands, providing a representational space that facilitates task performance. This would imply greater differentiation between the task-relevant and irrelevant stimuli neural patterns during attentional selection and reduced representational separation between stimuli in the integration task since both must remain relevant to the final decision and their information must be jointly considered.

## 2. Methods

### 2.1. Participants

We recruited forty-eight participants (mean age = 20.98; range = 18-27; 26 women and 22 men) from the University of Granada, all native Spanish speakers, right-handed and with normal or corrected vision. They were compensated with 30-35 euros, depending on their performance. Four additional participants were excluded due to low accuracy (below 80%) or more than 30% discarded EEG trials due to artifacts. All participants signed a consent form approved by the local Ethics Committee.

Sample size was calculated using PANGEA (Power ANalysis for GEneral ANOVA designs; Westfall, 2016). The experiment had a 3-factors within-subjects design (Task type x Congruency x Stimulus category) and our main contrast of interest was the two-way interaction Task type x Congruency. Attaining 80% statistical power for detecting a small-medium behavioral effect with a Cohen’s d value of 0.3 required a minimum of 30 participants. Due to counterbalancing needs we collected data from 48 participants, achieving 95% power.

### 2.2. Apparatus, stimuli and procedure

The task was run on Matlab 2020a using The Psychophysics Toolbox 3 (Brainard, 1997). Stimuli were presented on an LCD screen (1920 × 1080 resolution, 143,6 Hz refresh rate) over a grey background. To indicate the relevant category in each trial, we used 4 stimuli as cues, 2 per category: the letter A and a footprint symbol for the animal category, and the letter H and a gear symbol for the tool category. Target stimuli for each category were colored images of 8 animals (whale, chick, llama, wasp, crocodile, snail, bear, fish) and 8 tools (lawnmower, screwdriver, wheelbarrow, screw, cement mixer, knife, chainsaw, pliers). In each category, 50% were bigger than a shoe box and the rest smaller. In total, there were 512 images for cued animals trials and the same for tools. All stimulus images were retrieved from the Google image search engine using the CCBY 4.0 filter.

The experiment included 2 main tasks, selection and integration, and a separate stimulus localizer. Selection and integration tasks presented a cue followed by two stimuli (animal and tool) that required judging whether the size of the item in real life was larger or smaller than a shoe box. Cues indicated the category of the target (animal or tool) to pay attention to. In the selection task, participants had to judge the size of the cued target and ignore the additional stimulus. In integration, participants had to judge the size of the cued target and the additional stimulus, responding if both had the same size relationship with respect to a shoe box or not. Although in the integration condition the cue did not designate one stimulus as more task-relevant than the other, we kept the preparatory cueing information matched across tasks, providing a common basis for examining whether the cue-designated category was represented before target onset. The two stimuli could be congruent (i.e. same size relationship, 50%) or incongruent (i.e. different size relationship, 50%). Stimulus size relations and response categories were fully balanced across the two tasks. Within each stimulus category, half of the exemplars were larger than a shoebox and half were smaller. As all possible animal–tool combinations were presented, the integration task included equal numbers of congruent pairs (both stimuli larger or both smaller than a shoebox) and incongruent pairs (one stimulus larger and the other smaller). Consequently, the correct responses were balanced between “larger” and “smaller” in the selection task and between “same” (congruent) and “different” (incongruent) in the integration task, with each response category accounting for 50% of trials. At the beginning of each block participants were instructed about the task to perform (selection: “*in this block you have to pay attention only to the category of stimulus indicated by the cue to answer if this stimulus is bigger or smaller than a shoe box*”; integration: “*in this block you have to pay attention to the category of stimuli indicated by the cue and integrate it with the other stimulus to answer if they have the same size relationship with respect to a shoe box*”) and reminded the cue-category associations. Catch trials (20%) were included to ensure that cued categories were processed in the preparatory interval in both tasks (and not merely in selection trials, for a similar approach see e.g. Pena et al., 2025; González-García et al., 2021; Formica et al., 2021). In these, instead of the target and distractor, the words “ANIMAL” and “HERRAMIENTA” (animal and tool in Spanish; 10.47° x 0.67° degrees of visual angle, dva) appeared at the right and left side of the screen, and participants were instructed to respond with the left (key “D”) or right index (key “K”) to indicate the position of the word matching the cue’s category. The location of the words was randomized in each block. To prevent perceptual confounds in the multivariate analyses, 2 different cues were associated with each category (2 for animals, 2 for tools), and the 4 four cues were employed in each block. In localizer blocks the same animal and tool images were presented individually and participants had to press the key “C” for rotated stimuli (at 180°; 5.9% of the trials).

The sequence of events in each trial is displayed in Figure 1. The cue (∼ 2.86° x 2.86° dva) was shown during 50 ms and followed by a fixed Cue-Target Interval (CTI) of 1000 ms. Then an animal and a tool image (each ∼ 4.7° x 4.7° dva) appeared simultaneously for 200 ms. The two did not overlap. The response window started with target onset and lasted 1750 ms. The keys “D” or “K” were used to indicate the size relationship with a shoe box, with a S-R mapping fixed per participant and counterbalanced across them. The Inter-Trial-Interval (ITI) varied randomly between 2300 ms and 2800 ms. During the localizer trials, individual stimuli were presented for 200 ms followed by an ITI of 1000 ms.

**Figure 1.**
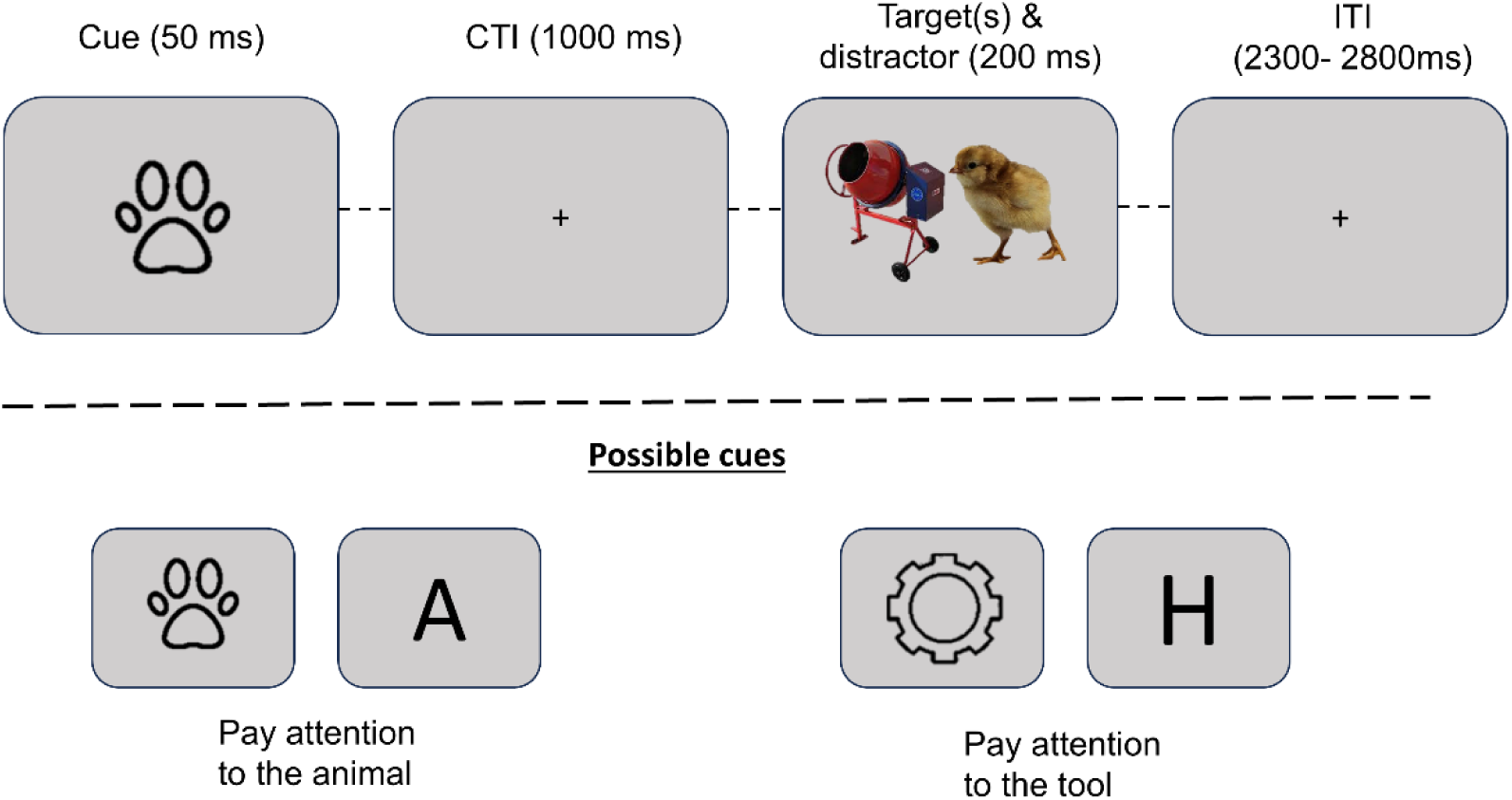
Experimental paradigm and example trial. In a size classification task, participants were cued about which stimulus category (animals or tools) they had to pay attention first. In selection blocks, participants had to respond only to the stimulus indicated by the cue (in the example, the animal is smaller than a shoe box), whereas in integration blocks they had to respond based on both stimuli (in the example, they don’t have the same size relationship with respect to a shoe box).

Participants read the instructions and were told the correct size judgement for each stimulus. Then, they performed a practice session where they had to achieve a minimum of 80% accuracy on each type of task, and more than 50% accuracy on catch trials. They completed a minimum of 2 practice blocks per type of task, repeating the full protocol up to 3 times until they reached the minimum accuracy. Selection, integration and localizer blocks were interspersed throughout the session with an order fully counterbalanced within and across participants, ensuring that each type of block was presented an equal number of times before and after the others. The task was composed of 48 blocks, 16 of each type (selection, integration and localizer). Each selection and integration block had 40 trials (8 catch). Each block lasted between 2.36 and 2.7 minutes (with 40 trials of 3.55 s – 4.05 s). The localizer blocks had 34 trials lasting 1.2 s, for a total of 40.8 s per block (544 localizer trials in total). Data from these localizer blocks were not used for the current study, so they will not be further discussed, as all analyses were based on trials from selection and integration demands. The whole session, including practice, lasted approximately 2 hours and 30 minutes. In total there were 512 trials (without catch) for selection and the same number for integration. These trials were unique combinations of the 8 animals and 8 tools, given that these 64 combinations could be preceded by the 4 different types of cues and the target could be presented on the right or left.

### 2.3. EEG acquisition and preprocessing

High-density EEG data was acquired using a 64-channel actiCap Slim (BrainVision) system at the Mind, Brain, and Behavior Research Center (CIMCYC) of the University of Granada. Impedances were kept below 10 kΩ, as recommended by the amplifier manufacturer. EEG signals were continuously referenced to the FCz electrode and digitized at a sampling rate of 1000 Hz.

Data preprocessing was performed using EEGLAB (Delorme & Makeig, 2004) and custom MATLAB scripts, following the pipeline available on GitHub (see Open Practices section). Data were downsampled to 256 Hz and a Finite Impulse Response (FIR) filter was applied to band-pass the signal between 0.1 and 126 Hz, plus notch filters at 50 and 100 Hz. Noisy channels were detected by visual inspection and removed to be interpolated at the end of the preprocessing (0.4 channels on average, range 0-3). Data was epoched in intervals of 3.5 sec (−1 to 2.5 seconds, locked to the cue). We applied ICA using the runica algorithm in EEGLAB to remove blink- and eye-movement–related components, identified by visual inspection and confirmed with ICLabel. On average, 1.3 components were removed per participant (range 1–3). Automatic trial rejection was applied using three criteria (see Aguado-López et al., (2024); López-García et al., (2022); Peñalver et al., (2023) for similar parameters). First, trials with abnormal spectral power, deviating from baseline by ±50 dB in the 0-2 Hz frequency range (sensitive to remaining eye artifacts) or by -100 dB or +25 dB in the 20-40 Hz range (sensitive to muscle activity) were excluded. Second, a probability distribution of voltage values across trials was computed and trials exceeding a threshold of ±6 standard deviations were rejected. Finally, any trial with voltage amplitudes ±150 mV in any electrode was excluded. Next, the removed channels were recomputed by spherical interpolation and a common average was used to re-reference the data. Finally, a baseline correction (−200 to 0 msec) was applied. Only correct non-catch trials were included in the EEG analyses, yielding an average of 873 trials per participant for the main task (range 736–969).

### 2.4. Analyses

#### 2.4.1. Behavioral

We performed 2-way repeated measures ANOVAs with the factors Task (Selection vs. Integration) and Congruency (Congruent vs. Incongruent) on accuracy and reaction times (RT) using the JASP software (Love et al., 2019). The factor stimulus category was not relevant for the behavioral analyses and was only considered in the EEG analyses. Catch trials were excluded. In RT data we excluded incorrect trials and those with RT deviating 2 standard deviations (SD) from the participants’ mean.

#### 2.4.2. EEG

##### 2.4.2.1. Multivariate pattern analysis (MVPA)

We employed Linear Discriminant Analysis (LDA) classifiers to analyze neural patterns focused on cue-locked activity from −100 to 2500 ms, conducted in MATLAB using the MVPAlab toolbox (López-García et al., 2022).

For each subject, classifiers were trained and tested on the raw voltage of each trial and time point across all channels, using identical configurations for all analyses. To enhance the signal-to-noise ratio, data were smoothed with a five-point moving average window. A 5-fold cross-validation scheme was applied to ensure unbiased performance estimation while controlling computational cost (Grootswagers et al., 2017). Data were subsampled so that both classes contained an equal number of trials, and class balance was maintained within each fold (Grootswagers et al., 2017; King & Dehaene, 2014). Normalization was performed within each fold by computing the mean (μ_train) and standard deviation (σ_train) of the training trials for each electrode, then applying these to normalize both training and test data:

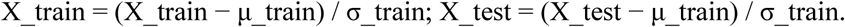

Analyses were computed every three time points to reduce computational load. Classification performance was quantified using the area under the curve (AUC), a non-parametric measure sensitive to binary differences and interpretable as classification accuracy (King et al., 2013; King & Dehaene, 2014).

The AUC values from all participants were averaged to create a group mean. Then, to assess group-level statistical significance, we applied a non-parametric cluster-based permutation test against empirical chance. Trial labels were randomly permuted 100 times per participant to generate null distributions, and group-level null decoding curves were computed by averaging one random AUC per participant. This process was repeated 10⁵ times to build empirical chance-level AUC distributions per time point. The 95th percentile of this distribution was used to identify significant decoding peaks. The minimum cluster size for significance (α = .05) was estimated from the permuted results and corrected for multiple comparisons using a False Discovery Rate (FDR) approach (López-García et al., 2022).

Additionally, we examined the extent to which activity patterns remained stable throughout the entire trial (King & Dehaene, 2014). To this end, we implemented a temporal generalization approach, applying the decoding analysis described above by training the classifier at a specific time point and testing it across all other time points. This procedure was iteratively repeated such that each time point served as both a training and a testing instance. The resulting output was a Temporal Generalization Matrix, in which the diagonal elements recapitulate the results of the classifier time course, while the off-diagonal elements capture the temporal generalization of the underlying neural representations. Statistical significance within these matrices was assessed using the same cluster-based permutation analysis described previously, extended here to identify two-dimensional clusters spanning both training and testing time points.

With this overall approach, we evaluated whether we could decode information about the target category, either animals or tools, during both the preparatory and target processing phases, performing classification analyses separately for selection and integration blocks. To reduce the possibility that category decoding reflected the physical appearance of the cues rather than category-related information, we used a cross-classification approach (Aguado-López et al., 2024; Kaplan et al., 2015; Peñalver et al., 2023). Specifically, classifiers were trained on trials preceded by one set of physical cues and tested on trials preceded by a different set of cues associated with the same categories, and vice versa. This approach required the classifiers to generalize across different cue identities, minimizing the influence of cue-specific visual features on category decoding. This protocol iterated across the two sides of the classification (exchanging training and testing cues) and results were averaged across directions.

Additionally, to test whether selection task increased the fidelity of category-specific neural codes, we compared the two cross-decoding curves and the temporal generalization matrices using a cluster-based permutation test. Specifically, for each subject and timepoint, we computed the difference between the two classification curves (or matrices). We repeated the same procedure after permuting the data labels, generating 100 difference curves per participant. From these, one curve per participant was randomly selected, averaged across subjects, and the process was repeated 10⁵ times to build a null distribution. Significant clusters were then identified using the same procedure described above.

##### 2.4.2.2. Representational similarity analysis (RSA)

The extent to which neural activity patterns were structured according to task demands and stimuli type was evaluated through RSA (Kriegeskorte et al., 2008). This analysis calculates the pairwise distances across all conditions to abstract the activity patterns into a common representational space. A time-resolved RSA was conducted to characterize the temporal dynamics of neural patterns across the entire trial epoch. The analysis followed similar configuration parameters to previous studies in our lab (Pena et al., 2025).

To construct the Representational Dissimilarity Matrices (RDMs), we considered all combinations of stimuli, cues and tasks. Specifically, the 16 possible targets (eight animals and eight tools) could be preceded by either of the two cue types (symbol or letter), yielding 32 possible stimulus–cue combinations. Each of these could appear in either of the two tasks, resulting in a total of 64 conditions (2 tasks x 2 types of cues x 16 targets) and, consequently, 64 × 64 matrices. Theoretical RDMs (Fig. 2A) were constructed to capture the expected distances between conditions under different hypotheses, with distances coded as 0 (minimum dissimilarity) or 1 (maximum dissimilarity). We specified five models based on our variables of interest: **task** (selection vs. integration), **category** (animal vs. tool), **exemplars** (8 animals, 8 tools), an interaction model of **task × category**, and an interaction model of **task × exemplars**. The three former models were built using 0s whenever two trial sets shared the corresponding task, category or exemplar conditions and 1s elsewhere, while the interaction ones were created multiplying the two original models’ matrices. As a complementary analysis, two additional models were included to account for potential perceptual biases driven by the cue identity and cue type (i.e., letter vs. symbol; see Supplementary Materials, Fig. S1). Note that the exemplar variable specifically assessed the neural coding of stimulus identity. Given that half of the exemplars were associated with the same motor response (left or right), motor response information on its own could not be used to generate above-chance exemplar classification.

**Figure 2.**
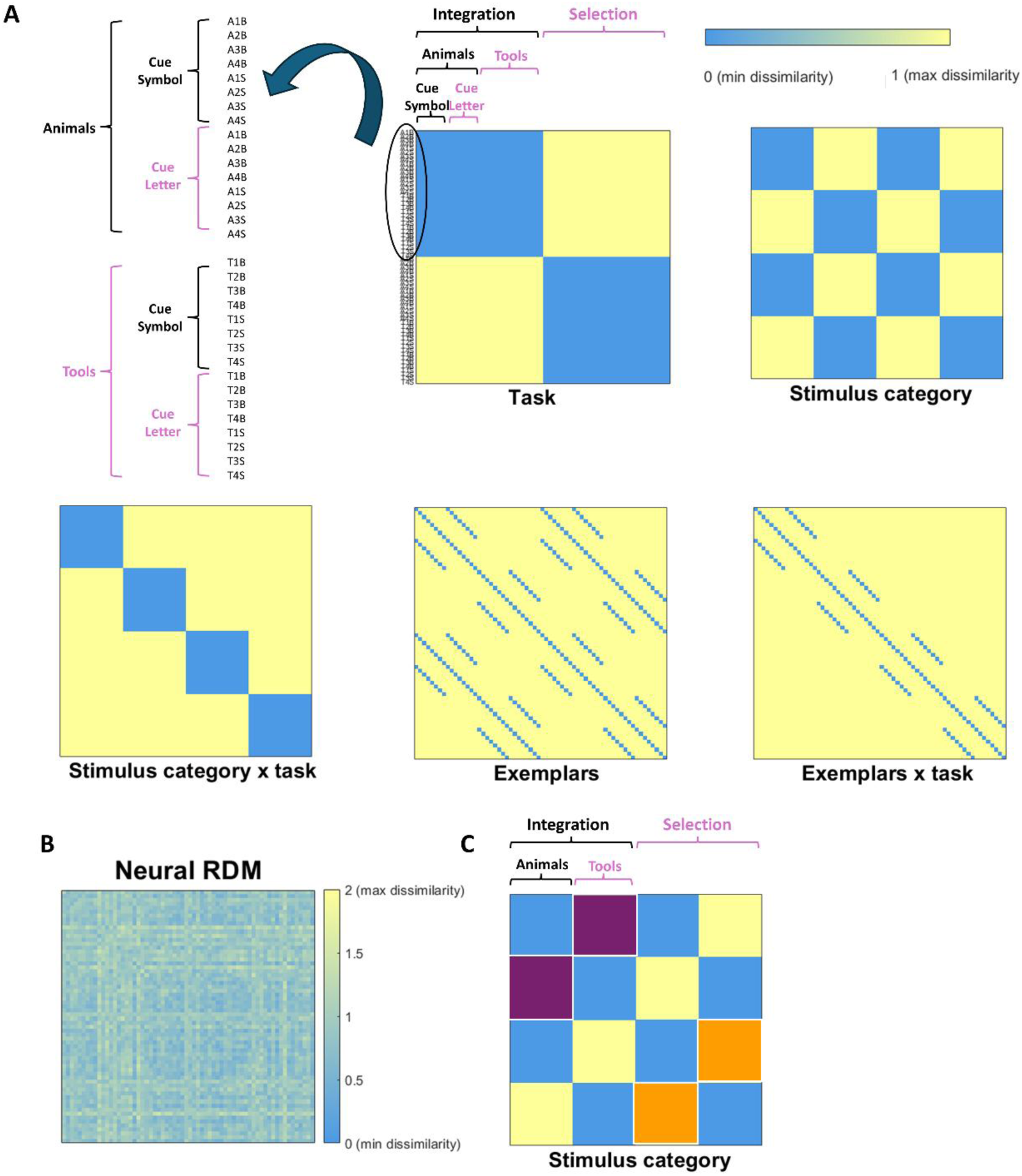
**A:** Model RSA matrices. The labels on the left indicate the cued stimulus in each task condition. A = animal; T = tool; B = big; S = small. **B:** Example of a neural RDM of one participant at one time point. **C.** Cells selected to average and compare the empirical dissimilarity curves, in orange for selection, in purple for integration.

Prior to the analysis, data were smoothed by applying a moving average window every five timepoints and normalized by z-scoring the values across all trials, regardless of the condition. For every condition and participant, EEG data were selected every three time points of the trial (−100 to 2500 ms locked to the cue) to reduce the processing load. The distance between each pair of conditions was estimated with a cross-validated version of Pearson’s correlation coefficient to reduce potential positive bias in the results (Grootswagers et al., 2017; Walther et al., 2016). For each participant we took all the data (timepoints and electrodes) and performed a threefold cross-validation procedure. We randomly split the data into three equally-sized chunks and, in each fold, one chunk served as testing set, while the remaining two were averaged and used as training sets. At every time point, we calculated the Pearson’s correlation for every pair of conditions between the training and test sets. The result was transformed into a distance metric by computing 1 - Pearson’s coefficient (i.e., 2 indicates maximum dissimilarity and 0 minimum dissimilarity). This process was repeated across all folds, ensuring each chunk served as test once. The resulting distance values were averaged across folds, yielding an empirical 64 × 64 RDM per participant and time point.

To estimate the share of variance explained by each theoretical RDM, we used multiple linear regressions. At each time point, we predicted the empirical RDM using the theoretical RDMs as regressors. Since cross-validated distance metrics were employed, we retained the full RDM and the diagonal in these analyses (Walther et al., 2016). All matrices were vectorized before performing the regressions. This procedure provided a beta coefficient and its corresponding *t-*value for each theoretical RDM, time point and participant, conveying the unique portion of the variance explained by the model.

To assess statistical significance at the group level, we focused on the regressions’ *t*-values and used a non-parametric cluster-based permutation method against empirical chance levels (López-García et al., 2022; Maris & Oostenveld, 2007). First, at the individual level, the values of the empirical RDMs were randomly shuffled and these values were used for a regression with the theoretical matrices. This process was repeated 100 times, creating 100 chance-level *t*-values per theoretical model and participant. Subsequently, one iteration was randomly selected for each subject, such that a single *t*-value per model was retained and then averaged at the group level. We repeated this process 10^5^ times to obtain 10^5^ permuted group *t*-values. These permuted maps were used to estimate the upper and lower bounds of the empirical chance distribution. Specifically, the 95th percentile was calculated at each time point, along with the 95th percentile of the cluster size of consecutive time points. To identify the minimum cluster size considered statistically significant at an alpha level of 0.05, we applied a correction for multiple comparisons using the false discovery rate (FDR).

To decompose the results of the interaction models (task * category, task * exemplar) we performed two additional RSAs analyzing, separately, selection and integration data. Following the same model-based approach described above, we computed the variance explained by the category and exemplar RDM models within each task condition. We then compared the resulting RSA curves (t-values) between tasks using a cluster-based permutation test equivalent to one employed to compare the two cross-decoding curves.

### Empirical RDM analysis

To determine whether task demands altered the representational separation between the two simultaneously presented stimuli, we extracted the empirical representational dissimilarity values. To do this, we focused on the estimated dissimilarity between pairs of stimuli of different categories (which were either target plus distractor under selection demands, or to-be-integrated stimuli), separately for each task, and averaged the corresponding RMD cells (see Fig. 2B) in each time-point. The resulting values formed two dissimilarity curves, corresponding to the dissimilarity between those stimuli in selection and integration task. These were compared with t-tests and then we identified cluster sizes by looking for sets of temporally adjacent points with p < 0.05. For each cluster, we computed a *t*-value by summing the *t*-values across all included time points. Statistical inference was then conducted by comparing these cluster-level values with a permutation-based null distribution. Specifically, for each permutation, condition labels were switched for half participants and repeated the same clustering procedure, generating 5,000 permuted cluster *t*-values. The resulting distribution was used to determine significance, with the 95th percentile serving as the threshold for identifying clusters in the empirical data as statistically significant (for a similar method see Aguado-López et al., 2024; Peñalver et al., 2023).

## 3. Results

### 3.1. Behavioral

Average accuracy was 91.15% (SD = 6.20%) for the main task and 93.33% (SD = 4.90%) for catch trials (94.40% catch in selection, 92.20% in integration). The ANOVA revealed a main effect of Congruency (F_47,1_ = 8.535, *p* = 0.005, *η*_p_^2^ = 0.154), indicating higher accuracy in congruent (M = 91.7%, SD = 5.8%) than incongruent trials (M = 90.6%, SD = 6.7%). There was also a significant interaction of Task * Congruency. T-tests showed that incongruent trials were performed worse than congruent in the Selection task (M _Cong_ = 93.1%, M _Incong_ = 89.0%, t _47,1_ = 7.535, *p* < 0.001, Cohen’s d = 1.088), whereas the opposite was found for the Integration task (M _Cong_ = 90.4%, M _Incong_ = 92.3%, t _47,1_ = -2.834, *p* = 0.007, Cohen’s d = -0.409, see Fig. 3).

**Figure 3.**
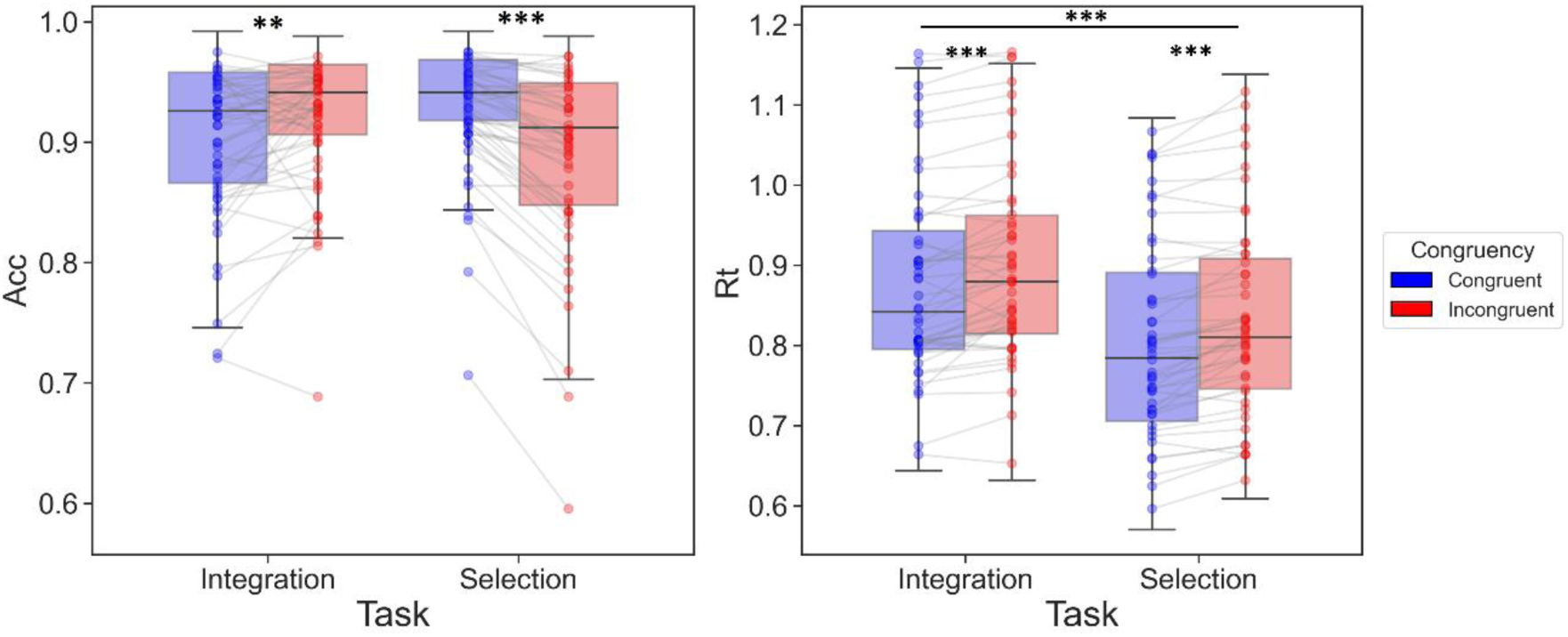
Box plots displaying the behavioral results. Boxes have a middle marking the median, limits representing the first and third quartile, and whiskers indicating the 1.5 inter quartile range for the upper and lower quartiles. Outliers are shown outside the whiskers. The dots represent each participant’s average per experimental condition. (Left) Behavioral accuracy rate (ACC) in Selection and Integration blocks for Congruent and Incongruent trials. (Right) Reaction times (RT, in seconds) in Selection and Integration blocks for Congruent and Incongruent trials. ∗∗ = *p* < .01; ∗∗∗ = *p* < .001.

Regarding RT data, an average of 87 trials (4.78%) per participant were excluded after filtering outlier trials (± 2SD from the participant mean). Average RT was 854 ms (SD = 127) in the main task, and 668 ms (SD = 88) for catch trials (656 ms catch in selection, 680 integration). The ANOVA showed a main effect of Task (F_47,1_ = 72.041, *p* < 0.001, *η*_p_^2^ = 0.605), since participants were faster in Selection (M = 816 ms, SD = 131) than in Integration (M = 891 ms, SD = 131). Also, there was an effect of Congruency (F_47,1_ = 54.646, *p* < 0.001, *η*_p_^2^ = 0.538) with overall faster congruent (M = 841 ms, SD = 128) than incongruent trials (M = 866 ms, SD = 127). The interaction effect was not significant (F_47,1_ = 0.253, *p* = 0.618, *η*_p_^2^ = 0.005, see Fig. 3).

### 3.2. Electrophysiology

#### 3.2.1. Category-specific coding

Classifiers trained and tested to discriminate the category template that was indicated by the cue (animals vs tools) indicated a significant effect of category (see Fig. 4 top). In the selection task, we found significant decoding clusters from the cue onset and during most of the window (86-133ms, 203-238ms, 320-367ms, 754-777ms, 836-871ms, 953-976ms, 1210-1234ms, 1351-1422ms, 1445-1586ms, 1633-1680ms, 1867-1902ms, 2043-2066ms), while in the integration task, we could only decode the category at the beginning of the preparation interval (86-121ms, 391-437ms, 484-519ms). When we compared these two curves, we found significant differences in clusters located 300ms before the target, during target presentation and 400ms afterwards (754-789ms, 1070-1172ms, 1199-1246ms, 1644-1680ms) where category coding accuracy was higher for selection than integration task.

**Figure 4.**
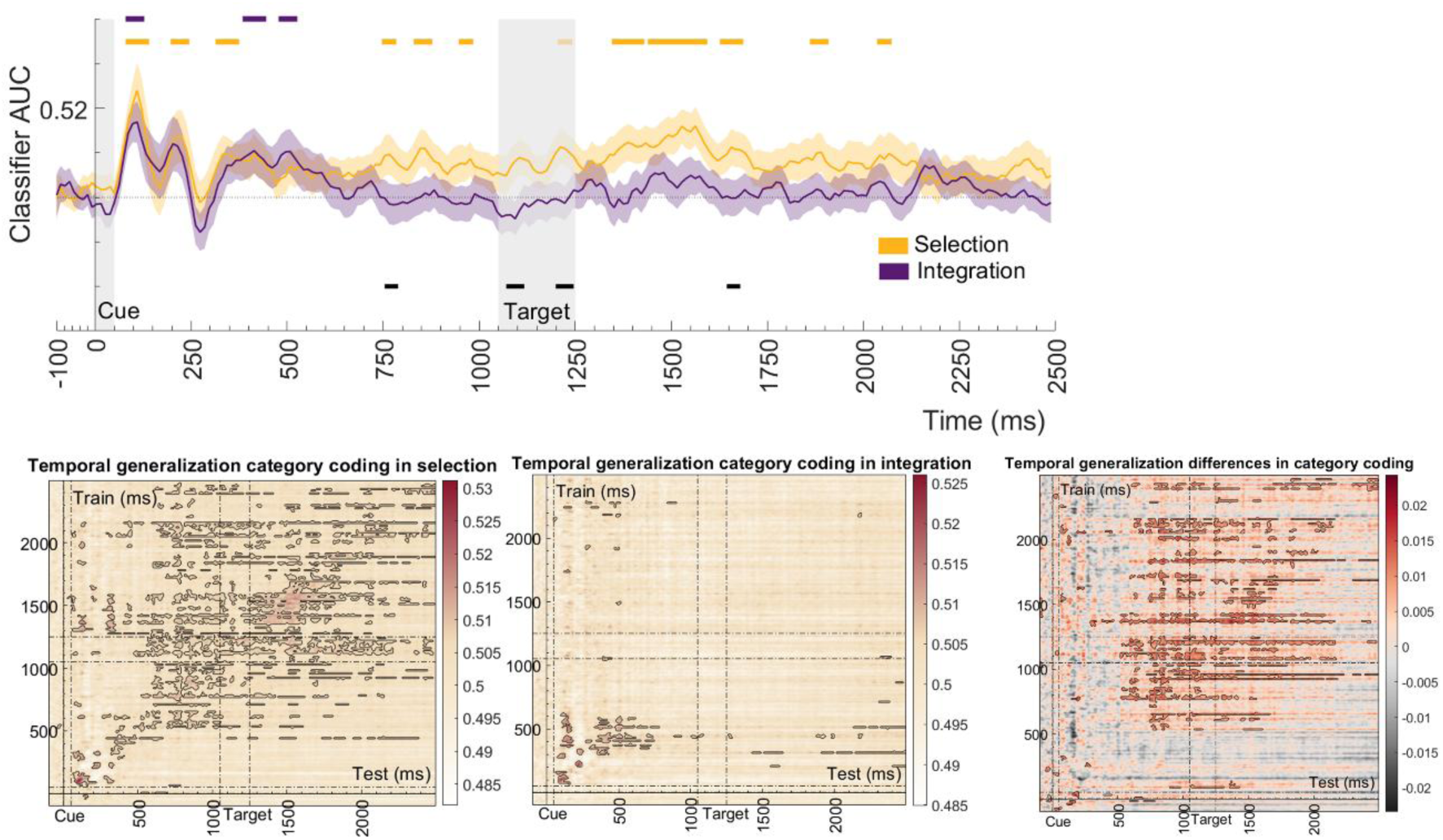
Top: Results (AUC values) of the classifiers discriminating the cued category (animals vs tools) using cross-classification (across cues identities), separately for selection (orange) and integration (purple) tasks. Colored shading around the lines represents standard errors of the mean (SEM). Gray shading indicates cue and target presence onscreen. Colored horizontal lines represent the significant clusters for selection and integration tasks, and black lines the significant differences between the two curves. Bottom: Temporal generalization matrices of the cued category coding. Significant clusters above chance are outlined with black. The color range in the bar represents AUC values in selection and integration. The subtraction differences between selection and integration are outlined in the last matrix. The cue and target presence onscreen is outlined with dotted lines.

The temporal generalization matrices of category-specific information showed different effects in selection and integration contexts (see Fig. 4 bottom). In selection, patterns generalized in the second part of the preparatory interval (after 500 ms), being more stable after target presentation (after 1050 ms). In integration, there were only a few clusters at the beginning of the preparation window. The comparison of the two matrices revealed statistically significant differences, corroborating the observed divergence in their temporal generalization patterns.

#### 3.2.2. RSA results

Using multivariate model-based RSA (Kriegeskorte et al., 2008), we compared neural data with the five RDMs of our variables of interest to assess how well each theoretical model captured the neural coding space across the trial epoch. The geometrical configuration of the activity patterns revealed that task demands were coded during a section of the preparatory period and became more stable after target presentation. Regarding stimulus coding, the neural geometry was significantly structured according to stimulus category during an early portion of the preparatory interval. This category-related RSA effect was strongest immediately following cue onset and decreased toward the end of the preparatory interval. On the other hand, specific exemplar configuration appeared during target presentation and the subsequent response period. Importantly, exemplars interacted with task demands progressively at the end of the trial, when response was needed (Figure 5A). Complementary analysis adding theoretical perceptual matrices showed similar results: all the variables mentioned remained significant. However, the effect of stimulus category was confined to a more limited time window and these perceptual matrices were also significant at the beginning of the preparatory interval (see Supplementary Fig. S2).

**Figure 5.**
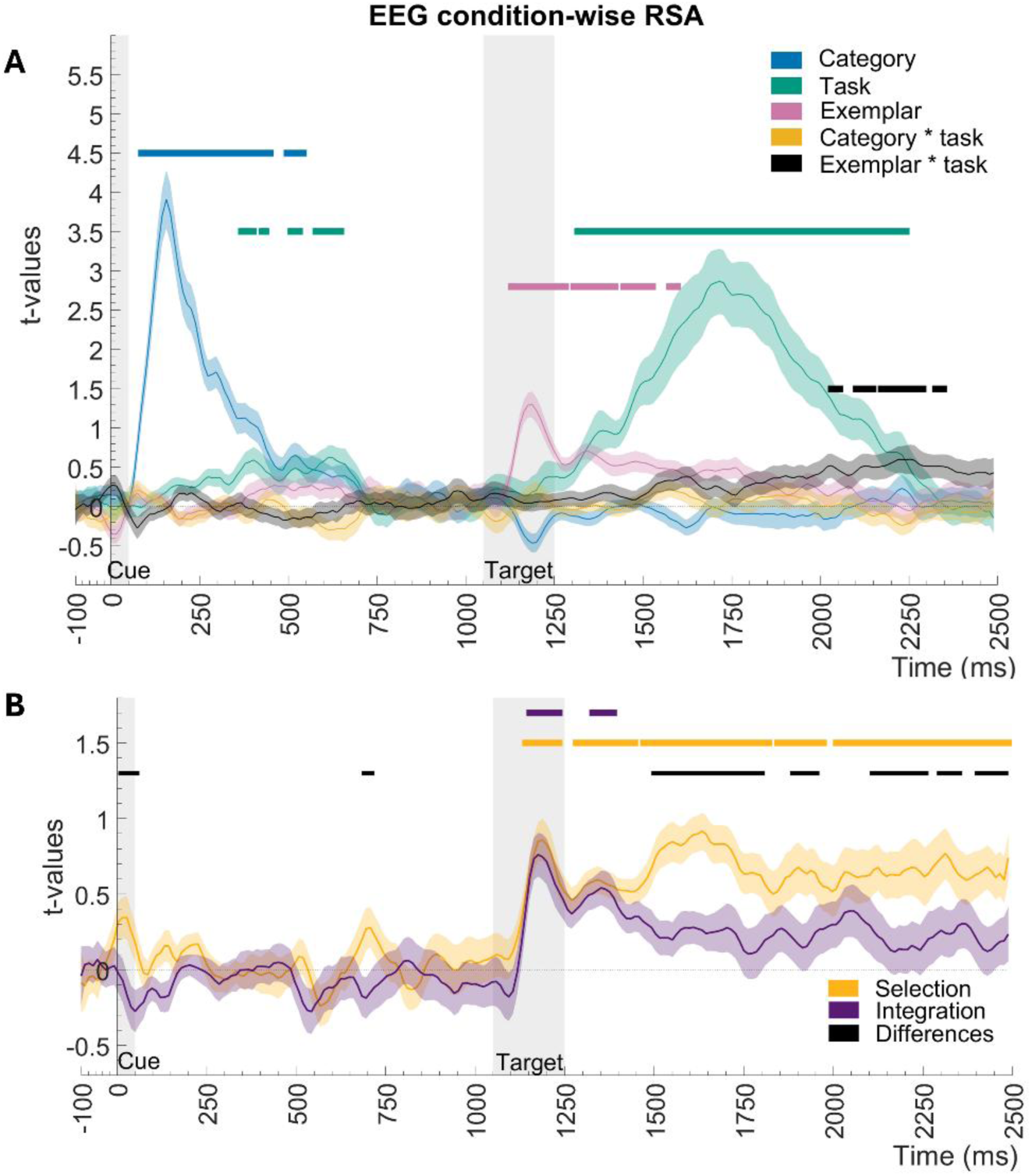
RSA time-resolved results. **A.** T-values after fitting the five theoretical models with a multiple linear regression, illustrating the unique share of variance explained by each of them. Each color represents a model, lighter shading refers to the standard error associated with the mean (SEM) t values. Gray shading indicates cue and target presence onscreen. The horizontal-colored lines above refer to the statistical significance against zero (dotted line) after implementing cluster-based permutation analysis. **B.** RSA time-resolved results of the exemplars matrix model, separately for selection (orange) and integration (purple) data. Gray shading indicates cue and target presence onscreen. Horizontal lines represent the significant clusters for selection and integration tasks, black lines represent the significant differences between curves. Note that exemplars where presented during target window, therefore there was no encoding before it.

Given the significant impact of the interaction model that predicted different geometries for exemplar coding under the two task demands, we performed additional analyses to better understand this phenomenon. The results showed that in the selection task, exemplars were encoded from stimuli presentation until the response is made, while in integration this coding was more transient and locked to target presentation (Figure 5B). Further analyses contrasting category encoding between tasks did not find significant differences (see Supplementary materials).

### Empirical RDM results

Finally, to determine whether selection and integration demands differed in the representational separation of the two simultaneously presented stimuli, we compared the dissimilarity values between stimulus categories in selection and integration tasks. This analysis is more fine-grained than model-based RSA since it reflects a direct measure of dissimilarity. Note that in this analysis we examined stimulus category and not exemplars, therefore dissimilarity values between categories could arise in the preparatory interval. Time-resolved results showed that representations conveying the two stimulus categories were more dissimilar in integration than selection after the cue and around the target. However, and in line with our hypothesis, categories became more dissimilar in selection than integration after target presentation, from 1633 ms until the end of the interval (Figure 6).

**Figure 6.**
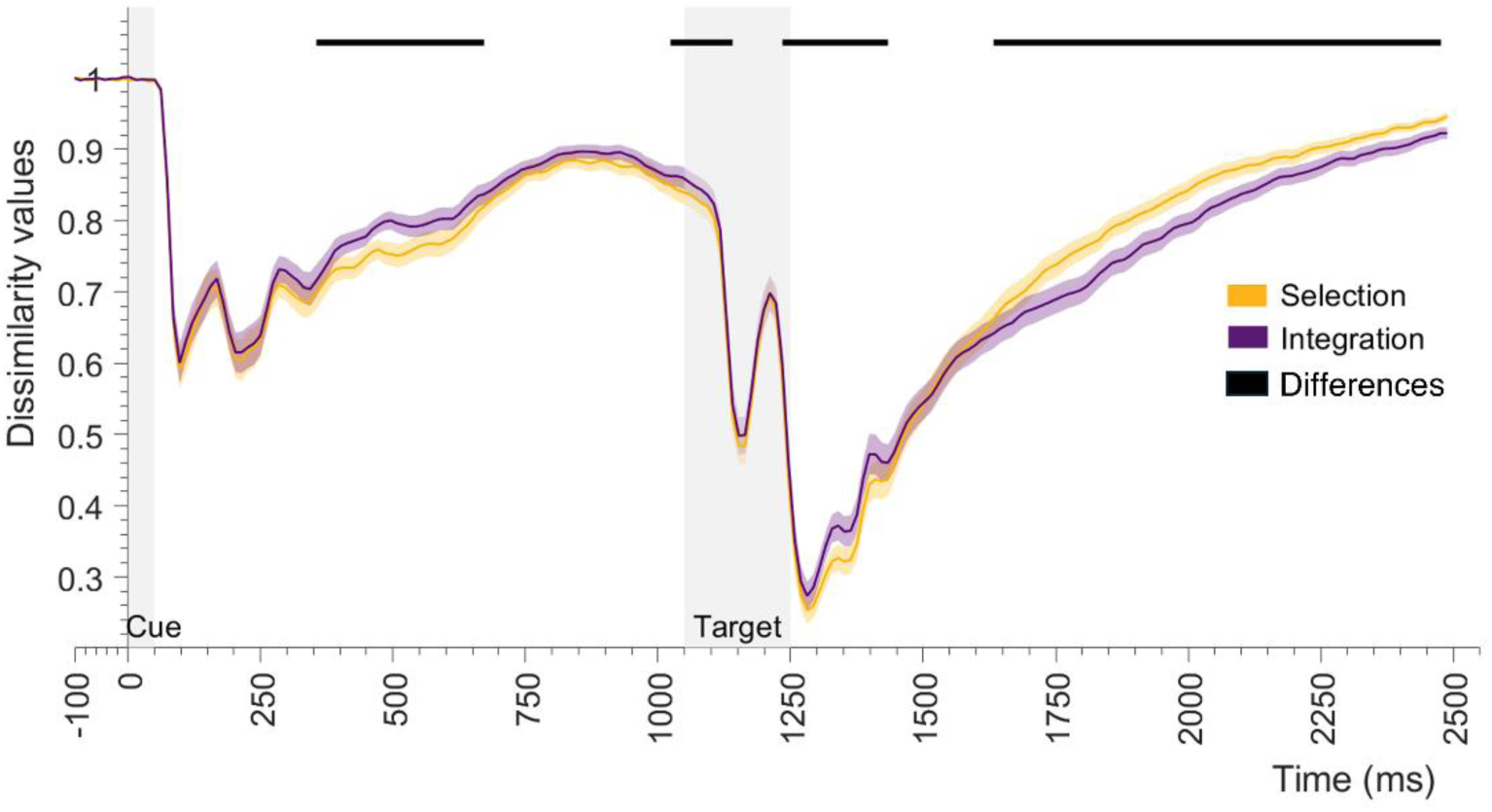
Empirical dissimilarity values between stimulus categories (cross-validated pearson values, from 0 to 2), averaged across different category cells separately for selection (orange) and integration (purple) tasks. Gray shading indicates cue and target presence onscreen. Black lines represent the significant cluster differences between curves. Note that these are raw dissimilarity values, and their direction differs from that of the multiple regression used in model-based RSA.

## 4. Discussion

In this study we investigated whether the attentional demands of selecting versus integrating information shape the brain coding geometry, during both preparatory periods and target processing. Using decoding analyses, we found that categorical stimulus templates were activated during the preparatory interval in both selection and integration contexts, whereas during actual stimulus perception they were only detectable under selection demands. Moreover, coding geometry assessed with RSA showed that stimulus coding was specific to exemplars at target perception, while during the subsequent response period, exemplars were encoded differently for selection and integration. Specifically, they were coded throughout the entire period following stimulus presentation under selection demands, whereas under integration demands they were coded only during target presentation. Further analyses indicated that during preparation and actual stimulus perception, dissimilarity between stimulus categories was greater in integration than in the selection, while the pattern reversed in the last part of the trial.

Our behavioral results confirm that participants were paying attention to the cues, as observed in the high accuracy rates in both task conditions and catch trials. Selection was performed faster than integration, which is coherent since integration requires considering more information (Fagioli & Macaluso, 2016; Morawetz et al., 2007). Moreover, in selection, the target congruency manipulation induced traditional congruency effects in both accuracy and RT, in line with previous studies and further validating our paradigm (Aguado-López et al., 2024; Desimone & Duncan, 1995; Egner, 2007; Eriksen & Eriksen, 1974). However, in integration blocks, congruent trials were less accurate than incongruent ones, suggesting that integrating information from similar real-word size stimuli is more laborious.

To our knowledge, this is the first study to compare the temporal evolution of multivariate neural coding during tasks that demand the selection or integration of information related to identity-based stimulus features. Consistent with our hypothesis, multivariate classifiers showed that the neural templates recruited by each task were tailored to its computational demands. While category-specific patterns were active for both tasks during the preparatory interval, in line with previous studies (Aguado-López et al., 2024; Esterman & Yantis, 2010; González-García et al., 2017; Peelen & Kastner, 2011; Peñalver et al., 2023), we found differences during target processing. Specifically, in the selection task classifiers revealed stable category coding across the preparatory interval, target processing and response period, consistent with prior findings in attentional selection paradigms (Cavanagh et al., 2018; Stokes et al., 2009). In contrast, in the integration task, category templates were detectable only during the preparatory interval. Importantly, although the role of the cue differed between tasks—participants judged only the cued stimulus in selection trials, whereas they judged both stimuli in integration trials—the presence of cue-related category information suggests that information associated with the upcoming stimulus set could be represented prospectively before target onset in both tasks. This temporal pattern aligns with our hypothesis: it suggests that in integration, the neural representation related to the cued category was no longer differentiable at the target processing, consistent with the possibility that processing became distributed across both stimuli representations rather than being preferentially organized around the cue-designated category. Importantly, we provided evidence supporting that the cued category templates were more separable at target presentation in selection compared to integration. This finding is consistent with the biased competition model (Desimone & Duncan, 1995), and could reflect the increased representational separation between categories in the selection task, but reduced separation when both stimuli must be considered. A complementary interpretation is that the presence of catch trials may have shaped participants’ strategies across tasks. Because catch trials were included in both conditions, participants may have encoded the cued (selected) category with greater fidelity during the preparatory interval, regardless of the selection or integration demands. Critically, although such a general up-regulation of attentional processes could have acted on top of demand-specific activity patterns, we believe it would not rule out our primary interpretation. While in the integration blocks the decisive moment may occur during target processing, when participants must determine whether to select a single category (as in catch trials) or integrate both categories (as in standard trials), this could happen already during the preparation in the selection task. This strategic flexibility could explain the differences found along the trial epoch between the integration and selection task. Additionally, the temporal generalization matrices indicated that, in the selection condition, category-specific patterns generalized from the late phase of the preparatory interval to the window of target processing, suggesting similarity between preparatory and target-related representational formats. Moreover, target-related coding remained stable throughout its processing period. This pattern is consistent with the notion of a preparatory template resembling the attentional representation of the selected target, as reported in prior work (Peñalver et al., 2024; Stokes et al., 2009). In contrast, this pattern was not observed in the integration task, where representations of the cued category generalized only during the early phase of the preparatory interval, consistent with our hypothesis.

Using RSA, we examined which dimensions structured the geometry of neural patterns and how they unfolded over time. We found that activity patterns encoded task information (i.e., selection or integration demands) during both the preparatory interval and actual target processing. This pattern aligns with previous work demonstrating task-specific neural patterns encoded anticipatorily (Aguado-López et al., 2024; González-García et al., 2017; Hall-McMaster et al., 2019; Manelis & Reder, 2015; Palenciano et al., 2019) as well as up until response execution (Pena et al., 2025). Still, the activity patterns coding task information in our data were not sustained throughout the entire preparatory interval. Instead, significant coding emerged only midway through preparation to later reappear during the response period. This result differs from earlier findings of sustained profile of preparatory task coding (Aguado-López et al., 2024; Pena et al., 2025; Stokes et al., 2013). One possibility is that, after encoding the cue, participants briefly store the task in an “activity-silent” state, minimizing active neural maintenance, until the task became relevant again at target onset. Consistent with this interpretation, we did observe a ramping-up of task representation before target presentation and its reactivation during the response window, in line with theories of dynamic coding and activity-silent working memory (Stokes, 2015).

Consistently, our RSA results also support the presence of categorical stimulus templates at the early part of the preparatory interval. The transient nature of this effect indicates that at the latter part of preparation, the neural activity patterns became less strongly structured according to the categorical geometry captured by this model, which may reflect a change in the representational format of the cued information. However, it does not necessarily imply that the cued category was no longer represented. This interpretation is supported by our complementary decoding analysis, where we found that category was also coded at the end of the preparatory interval for the selection condition. Therefore, category information can remain available while its representational geometry is not continuously captured by the category RSA model. Importantly, the RSA analysis also revealed more fine-grained stimulus representations detectable in neural pattern geometry. Accordingly, activity patterns coding the specific exemplars within the cued category were activated during target presentation and maintained thereafter, suggesting that specific rather than categorical neural codes govern target processing and response selection stages. Such exemplar-specific representations are consistent with previous findings (Carlson et al., 2013; Cichy et al., 2014; Kriegeskorte et al., 2008). More importantly, and as expected, fine-grained stimulus coding was tailored to task demands at the response interval, where we found that exemplar representations were more stable in the selection task while for the integration they were only detectable at target presentation. This is in line with the temporal differences we found in the categorical templates using classifiers, which showed sustained category information during target processing and the response interval in the selection condition, but weaker category decoding in the integration condition, potentially reflecting the different role of the cue in this task. Nevertheless, RSA revealed exemplar-specific representations during target processing, suggesting that reduced category-level decoding did not necessarily reflect an absence of stimulus information, but rather differences in the representational structure of the neural patterns. Overall, our results are consistent with the biased competition model. In the selection task, the task-relevant exemplar is prioritized and the competing stimulus is attenuated, thereby enhancing its representational distinctiveness. In contrast, when attention is devoted to integration, both exemplars become task-relevant, reducing the competitive advantage of the cued stimulus. Note that each exemplar had a fixed real-world size relation, meaning that exemplar identity could be partially confounded with the required response. However, the RSA specifically assessed the neural coding of stimulus identity, whereas the same motor response was associated with half of the exemplars. We therefore interpret the exemplar RSA results primarily as evidence for task-dependent stimulus-related representational structure.

Based on these task requirements, we hypothesized that the estimated neural similarity between exemplars from different categories would be greater in integration than in selection, reflecting the need to attend to and combine information from both categories. In line with this prediction, further analyses of dissimilarity values showed that in the response window, activity patterns from trials with different cued categories had a reduced separation in the integration than in the selection task. This pattern suggests that different categories were flexibly encoded depending on the overarching demands, converging onto a shared neural coding sub-space when both had to be jointly processed, or increasing their representational distance when selection (and competition) was needed. These dissimilarity effects emerged during the final portion of the trial. Notably, our analyses encompassed the entire response window; therefore, depending on the trial, these differences may have occurred either before or after the response. Even so, relative to the mean reaction time, reduced separation between stimuli in integration was observed prior to the response. It is worth noting that the direction of the effect was reversed in the preparatory and target processing interval, where reduced separation between categories was found in the selection task. One possible explanation is that, during preparation and early target processing of the selection task, participants maintain both categories active (i.e., cued and to-be-ignored categories) as a strategy to keep track of which category must be selected and which ignored before competitive suppression fully unfolds. This transient coexistence could reduce representational separation early in the trial epoch, which later shifted to the opposite -and hypothesized-pattern, with increased distance for selection demands. A non-exclusive alternative is that these representational dynamics reflect computations of different nature performed in specific brain areas, that were indistinctively reflected in our EEG patterns. Hence, the current findings would benefit from complementary spatially resolved neuroimaging techniques to identify the anatomical substrates underpinning preparatory and target-related dynamics in integration tasks. Fusion analysis of EEG and fMRI data (Cichy & Oliva, 2020) may help localize the regions supporting these temporally resolved neural processes. Finally, future studies using stimulus-pair matrices and larger trial counts are needed to further characterize this phenomenon, and could additionally employ trial-wise RSA approaches to test hypotheses about conjunctive representations (Kikumoto & Mayr, 2020).

It is important to highlight that we carried out a complementary RSA analysis showing that at the preparatory interval there was perceptual information about the cue. Although stimulus category encoding was still significant after removing the variance corresponding to the perceptual part of the cues, it became less prominent, suggesting that there were overlapping effects. Further studies counterbalancing the meaning of the cues across participants would be key to extend our results. Lastly, given the novelty of the paradigm, additional work is needed to examine other forms of integration, for example, tasks that require comparing the two stimuli directly or performing alternative cognitive operations across stimulus features, rather than comparing each stimulus independently before integrating them.

In conclusion, this study advances our understanding of the neural representational dynamics underlying attentional selection and integration demands. While preparatory category templates were found in both tasks, these differed in the neural representations recruited during target processing, where we found more sustained and robust exemplar coding in the selection than in the integration condition. Furthermore, task demands systematically shaped the similarity structure of stimulus representations in a goal-oriented manner: increasing the representational separation between the task-relevant and irrelevant stimuli during selection, while reducing their separation when both stimuli contributed to the task. Integrating these findings within a unified theoretical framework of attention will be essential for developing a more comprehensive account of top-down control mechanisms.

## Open practices

The data and materials used in this study are publicly available. Original code for the EEG preprocessing has been posted on our team Github: https://github.com/Human-Neuroscience/eeg-preprocessing. Raw EEG data, organized following the BIDS format (Pernet et al., 2019) will be deposited in the OpenNeuro public server once the paper is published. The analysis code, results and task code are shared in an Open Science Framework repository that will be public once the paper is published.

No part of the study procedures or analyses was preregistered prior to the research being undertaken. We report how we determined prior to our analyses our sample size, all data exclusions, all exclusion criteria, all manipulations, and all measures in the study.

## CRediT authorship contribution statement

**Blanca Aguado-López:** Writing – original draft, Software, Methodology, Conceptualization, Formal analysis, Data curation. **Ana F. Palenciano:** Writing – review & editing, Supervision, Formal analysis, Methodology, Conceptualization. **María Ruz:** Writing – review & editing, Supervision, Resources, Project administration, Funding acquisition, Methodology, Conceptualization.

## Acknowledgments and fundings

This research was supported by grants PID2019-111187GB-100 and PID2022-138940NB awarded to M.R. and Grant PID2023-151911NA-I00 awarded to A.F.P, all funded by MICIU/AEI/10.13039/501100011033 and ERDF/EU. B.A.L. was supported by a scholarship from the Spanish Ministry of Universities (FPU20/01980). A.F.P. was supported by Grant PAIDI21_00207 of the Andalusian Autonomic Government. The Mind, Brain and Behavior Research Center receives funding from Grants CEX2023-001312-M by MCIN/AEI/10.13039/501100011033 and UCE-PP2023-11 by the University of Granada. This study was conducted as part of B.A.L.’s doctoral research. We are grateful to Marta Becerra Losada and Juan Manuel Quesada Soto for their assistance with data collection.

## Appendix A. Supplementary materials

**Figure S1.**
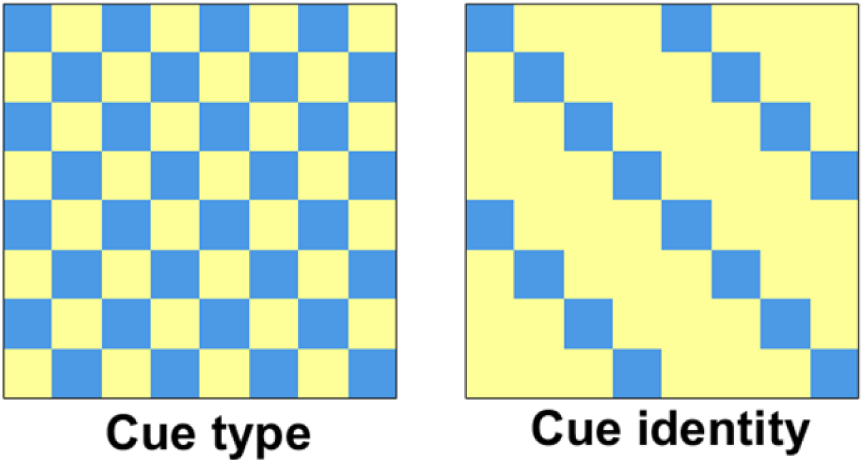
Additional model RDMs to control.

**Figure S2.**
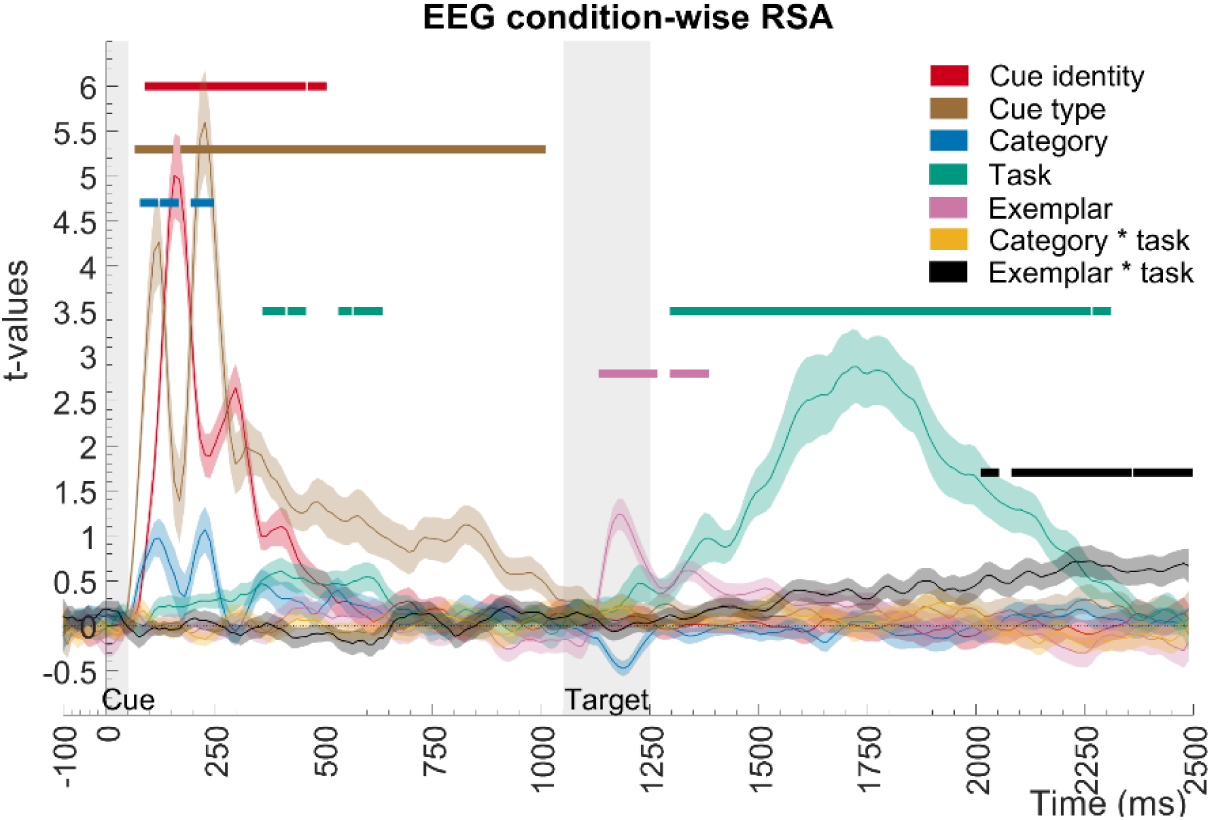
RSA time-resolved results adding 2 extra theoretical models: cue identity (4 different cues), cue type (symbols or letters). Each colored line represents the t values after fitting the seven theoretical models with a multiple linear regression, illustrating the unique share of variance explained by each model. Lighter shading refers to the standard error associated with the mean (SEM) t values. Gray shading indicates cue and target presence onscreen. The horizontal-colored lines above refer to the statistical significance against zero (dotted line) after implementing cluster-based permutation analysis.

**Figure S3.**
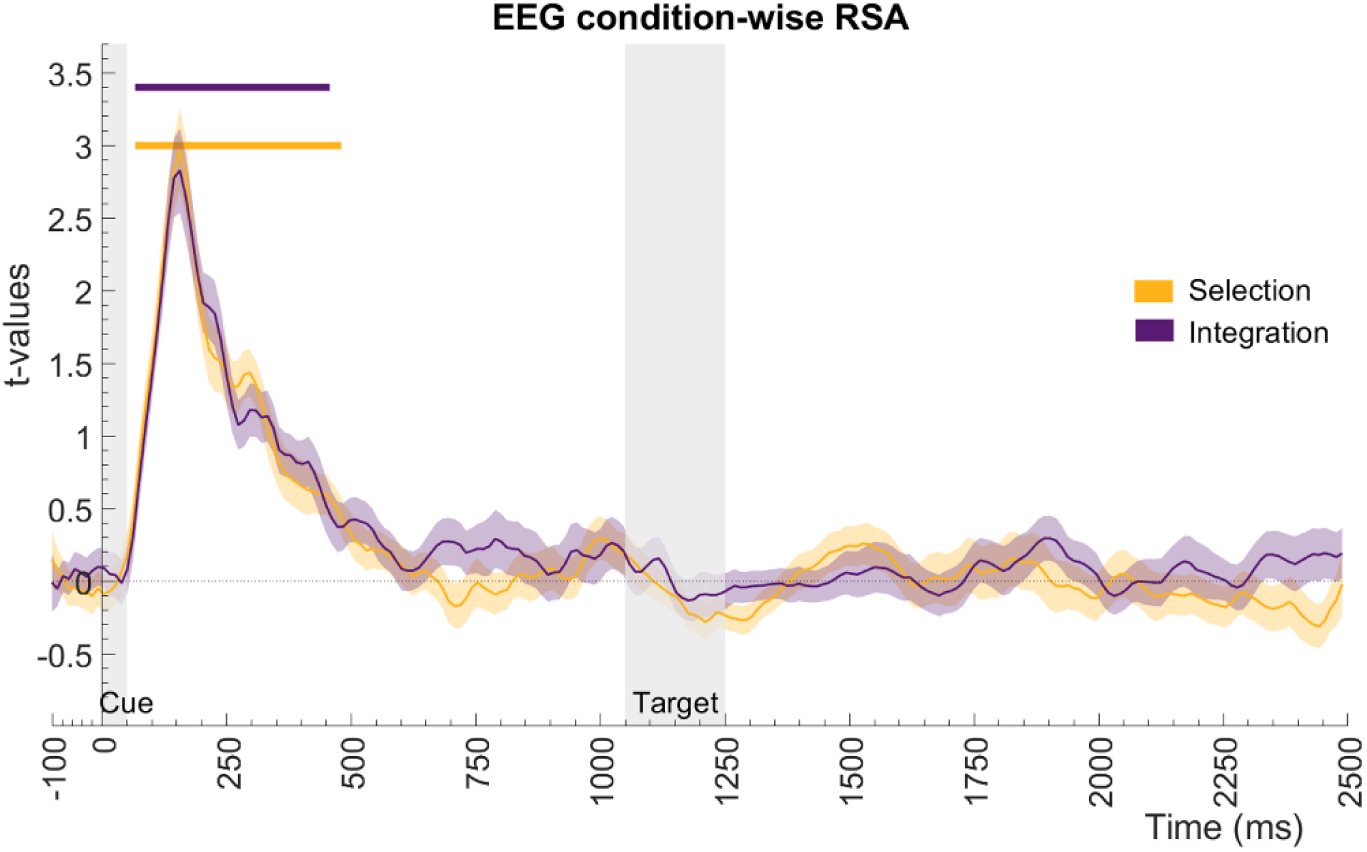
RSA time-resolved results of the stimulus category matrix model, separately for selection (orange) and integration (purple) data. Gray shading indicates cue and target presence onscreen. Horizontal lines represent significant clusters for selection and integration tasks.

